# Genome Mining and Pangenome Analysis of the *Stutzerimonas* Genus: a Novel Source of Plastic-Degrading Enzymes

**DOI:** 10.64898/2026.02.13.705723

**Authors:** Anna Luiza Bauer Canellas, Mauro de Medeiros Oliveira, Yeda Idalina Ilheo Rodrigues, Bruno Francesco Rodrigues de Oliveira, Nathália Ferreira dos Santos, Maria Alice Zarur Coelho, Johannes H. de Winde, Marinella Silva Laport

## Abstract

Nowadays, finding new sustainable ways to combat plastic pollution is a pressing challenge. Here, we provide a comprehensive genome mining analysis of 284 publicly available *Stutzerimonas* genomes for potential PET-active enzymes (PETases). While *Stutzerimonas* is a relatively newly established genus, it emerges as an interesting candidate in the search for novel biocatalysts. Hence, the first pangenome assessment of this genus based on its high-quality publicly available genomes was performed. An increasingly open pangenome was revealed, suggesting the versatility and adaptability of these strains to a variety of ecological niches. Moreover, functional characterisation of a new isolate, *Stutzerimonas frequens* VG-9, was carried out, confirming that enzymes found via *in silico* analyses may indeed display activity towards different polyesters. In summary, this study provides insights into the diversity of PETase homologues within still underexplored bacterial hosts, offering new perspectives for enzyme discovery in the Pseudomonadaceae family.

**Impact Statement:** Microbial enzymes known as PETases have emerged as promising candidates for the biological degradation of PET. This study investigated the potential of underexplored bacterial genera by genome mining of PETase homologues. Our findings provide new insights into the distribution of PETase-like enzymes in the Pseudomonadaceae family, offering a more comprehensive view of their plastic degradation capacity. These results hold practical implications for the development of optimized enzyme discovery strategies, while also highlighting the vast genetic plasticity of Pseudomonadaceae. We also provided the first report on the *Stutzerimonas* pangenome and insights into the enzymatic activity towards polyesters of a newly isolated strain. Hence, the role of this genus as a highly adaptable and versatile entity was reinforced, further disclosing it as a potential source of novel biocatalysts.

**Data Summary:** The genome of *S. frequens* VG9 has been deposited in Genbank under the accession number SAMN49487720. The accession numbers of all analyzed genomes are listed in Tables S2 and S3 (available in the online Supplementary Material).

## Introduction

The large-scale production and use of plastics dates to the 1950s (Geyer et al., 2017). In 2023 alone, global plastic production reached 436 million metric tons, of which 90.4% were estimated to be derived from fossil sources. The widespread use of these synthetic polymers across many fields and for varied purposes is immediately followed by more critical challenges, such as their proper disposal and potential repurposing. Indeed, up to 45% of the plastics produced are intended for single-use only, such as food packaging (Plastics Europe, 2024; Gates and Crook, 2024; United Nations, 2025). As a result of insufficient recycling policies and cost-effective and sustainable degradation strategies, a significant proportion of plastics is disposed of in natural environments, where they persist for long periods of time (Ali et al., 2021).

On an evolutionary timescale, the relatively recent large-scale introduction of plastics into natural ecosystems has not yielded sufficient time for the evolution of microorganisms specialised in the uptake of these polymers. One notable example of bacterial adaptation to plastic pollution is the isolation of *Ideonella sakaiensis* (now named *Piscinibacter sakaiensis*) from environmental samples of a polyethylene terephthalate (PET) bottle recycling site, a species that can degrade PET using two main enzymes, namely PETase and MHETase (Tanasupawat et al., 2016; Liu et al., 2022). However, such microorganisms are still relatively rare to find. Commonly, they appear adapted to the degradation of natural polymers - e.g. cutin, lignin - by repurposing their enzymatic machinery to target synthetic polymers (Gates and Crook, 2024). Hence, many of the known PET active enzymes, i.e. PETases are often annotated as amidases, cutinases, esterases, lipases or proteases (E.C. 3.1.x) (Bucholz et al., 2022).

In this context, the search for novel biocatalysts is a propitious venue of research in the field of plastic biodegradation (Verschoor et al., 2022; Qiu et al., 2024). These novel enzymes can be discovered by different strategies, such as *in vitro* screening with specific substrates (e.g. tributyrin, polycaprolactone) or *in silico* (meta)genome mining (Mollitor et al., 2019; Frey et al., 2024). Genome mining has been integrated as a robust approach into microbial bioprospecting efforts for enzyme biodiscovery (Ngyuen et al., 2024), being applied with success to reveal potential plastic-degrading enzymes by extensive large-scale analyses involving comparison of annotated genomic data using BLAST and profile hidden Markov models/HMMer (Dhali et al., 2024). *In silico* mining of plastic-degrading enzymes offers a time-effective and high-throughput screening approach to explore the enormous diversity of bacteria from varied environmental niches (Kim et al., 2022; Herbert et al., 2022; Zhu et al., 2022). A significant proportion of PETases are derived from bacteria, among which the family Pseudomonadaceae stands out as a source of many typical enzymes, including PE-H from *Pseudomonas* (*Halopseudomonas*) *aestusnigri* VGXO14^T^ (Bollinger et al., 2020), PpPETase from *Pseudomonas paralcaligenes* MRCP1333 (Sagong et al., 2021), and RgPETase from *Rhizobacter gummiphilus* NS21 (Han et al., 2024).

Hence, this study aimed to investigate whether the *Stutzerimonas* genus could be a useful target of PETase mining. We delved into the genomic and functional aspects of PET degradation by a newly isolated strain, *Stutzerimonas frequens* VG-9, and employed a custom PETase homologue database to further investigate high-quality and publicly available genomes. The findings provide novel and comprehensive insights into the distribution of PETase homologues across this genus, uncovering promising candidates for future functional studies. In light of the extremely large genetic diversity for biocatalytic discovery in this family, we extended our scope and embarked on reconstructing the pangenome of the *Stutzerimonas* genus, recently established after the promotion of the metabolically versatile *Pseudomonas stutzeri* species to a higher taxonomic rank (Lalucat et al., 2022). This is the first report of the *Stutzerimonas* pangenome, revealing unique traits pertaining to this relatively new genus and highlighting it as an interesting source of potentially novel biocatalysts.

## Materials and methods

### Functional screening of Stutzerimonas frequens VG-9

The strain *Stutzerimonas frequens* VG-9 was isolated from a water sample collected in Pedra Vermelha (22°59′07.8′′S 41°59′32.4′′W), an environmental protection area in Arraial do Cabo, Rio de Janeiro, Brazil. The strain was originally isolated in Glutamate Starch Phenol-red (GSP) agar and was submitted to enzyme activity screening on solid media (Luria Bertani) using multiple substrates, including: bis(2-hydroxyethyl) terephthalate (BHET 10 mM, Sigma Aldrich), Impranil DLN-SD (0.5%; Covestro), polycaprolactone (1%; PCL, Sigma Aldrich), tributyrin (1%; Sigma Aldrich), and tween 20 (1%) according to Canellas and colleagues (2023) and Carr and colleagues (2022).

### Degradation of BHET and MHET by S. frequens VG-9

*S. frequens* VG-9 was further evaluated for the ability to hydrolyse BHET and MHET in liquid culture. Cells of strain VG-9 pre-cultured in LB medium (24 h) were collected by centrifugation (4000 × *g*) for 10 min and inoculated in LB medium containing 500 mg.L^-1^ of BHET or mono-2-hydroxyethyl terephthalate (MHET) for an initial cell concentration of approximately 1 mg.L^-1^. All experiments were conducted at 28 °C and 250 rpm in an orbital incubator. Each experimental condition was performed in duplicate, with measurements taken daily after inoculation. Briefly, 1 mL of each sample was centrifuged (4000 × *g*) for 10 min and the supernatant was mixed with methanol (1:10) for further characterization.

The concentration of BHET or MHET and the formation of terephthalic acid (TPA) were detected by High Performance Liquid Chromatography (HPLC, Thermo Scientific, Dionex UltiMate 3000) using the Eclipse Plus C18 column, 5 μm, 4,6×250 mm, and the Zorbax SBC18, 5 μm, 4,6×125 mm. The injection volume of the sample was 10 μL, column temperature was maintained at 30 °C and the products were detected in a UV cell (254 nm). A gradient of acetonitrile and 0.05% (v/v) formic acid was used as the mobile phase, at a flow rate of 0.5 mL/min. The reference chromatograms of TPA, MHET, and BHET standards were used to identify BHET and its monomers. For each experiment, the parameters evaluated during the process of TPA production from BHET or MHET were the final concentration of TPA, yield factor (Y_p/s_), volumetric productivity (Q_p_) and bioconversion efficiency (BE). A metabolic stoichiometry of 1 mole of BHET or MHET per 1 mole of TPA was used to calculate bioconversion efficiency.

### Genome sequencing of S. frequens VG-9

An additional genome of a *S. frequens* strain (VG-9) sequenced by our research group was included in the analysis (Genbank accession number: SAMN49487720). Genomic DNA extraction was performed using the Wizard Genomic DNA Purification Kit A1120 (Promega, USA), following the manufacturer’s instructions. Library preparation was performed using the Nextera^®^ XT kit. The PCR products were purified using the AMPure^®^XP system and quantified using qPCR with the KAPA^®^ Fast Universal kit. Genomic DNA was sequenced using Next-Generation Sequencing (NGS) on the Illumina NextSeq500 platform. The quality of the read libraries was assessed using the FastQC (v0.11.9) tool (Andrews, 2022). Raw reads were processed using the Trimmomatic v0.39 software (Bolger et al., 2016), and the processed reads were used for genome assembly with SPAdes (v3.13.1) (Bankevich et al., 2012). The genome assembly was reoriented using the DNAapler tool (v0.7.0) (Bouras et al. 2024). The quality of the assembly was evaluated using assembly-stats (v1.0.1), seqkit (v2.1.0), CheckM (v1.2.3) (Parks et al., 2015) and BUSCO (v5.6.1) (pseudomonadales_odb12) (Tegenfeldt et al. 2025). The Average Nucleotide Identity (ANI) was calculated by comparing the genome to the reference genomes of *Stutzerimonas frequens* strain FDAARGOS_877 (GCF_016028515.1), TF18 (GCF_026639235.1), RL-XB02 (GCF_043636045.1), R2A2 (GCF_003205815.1), L27 (GCF_032770195.1), THAF7b (GCF_009363815.1), TPA3 (GCF_017577085.1), XN05-1 (GCF_035835015.1) and outgroup *Thermococcus kodakarensis* strain KOD1 (GCF_000009965.1) using the OrthoANI tool (Lee et al. 2016). Structural and functional annotation of the DNA sequences were performed using Bakta (v1.5.1) (Beyvers et al. 2025) and BlastKoala (Kanehisa et al. 2016). The identification of prophages within bacterial genomes was carried out using the PHASTEST (v3.0) (Wishart et al. 2023). The GTDB-Tool Kit (GTDB-Tk) was used to taxonomically classify the strain based on the presence of 120 single-copy marker genes in their draft assemblies and the placement of their genomes in the Genome Taxonomy Database (GTDB) reference tree using the available database (GTDB v2.4.1) (Chaumeil et al. 2019). To identify antibiotic resistance genes (ARGs), protein-coding nucleotide sequences from each genome were compared against different databases, including NCBI AMRFinderPlus (v3.0.12) (Feldgarden et al. 2021), CARD (v3.0.3) (Alcock et al. 2023), ResFinder (v4.0) (Bortolaia et al. 2020), ARG-ANNOT (v6) (Gupta et al. 2014), MEGARES (v2.0) (Doster et al. 2020), and PlasmidFinder (v2.0.1) (Carattoli et al. 2014) using the ABRICATE tool (v1.0.1) (https://github.com/tseemann/abricate). Additionally, the genome was analysed for the presence of Virulence-related genes using the ABRICATE tool and databases VFDB (v2019) (Chen et al. 2016). Biosynthetic gene clusters (BGCs) associated with antimicrobial activity were identified using antiSMASH (v8.0) (Blin et al. 2025), enabling the detection of additional regions linked to other secondary metabolites. The CLEAN web server was used to assign Enzyme Commission (EC) numbers to amino acid sequences (Yu et al., 2023). Putative plasmids were identified using two plasmid prediction tools: Deeplasmid (Andreopoulos et al. 2022) and Plasme (v1.1) (Tang et al. 2023) with a score above 0.9. Circular genome representations were generated using the GenoVi pipeline (v0.4.3) (Cumsille et al., 2023).

### PETase database construction and genome mining

A custom PETase database was constructed based on strategies previously described (Carr et al., 2022; Howard et al, 2022), with modifications. Here, 75 sequences were retrieved from the Plastics-Active Enzymes Database (PAZy), encompassing biochemically characterised PET-active enzymes (Buchholz et al., 2022) (**Table S1**). For *Stutzerimonas* (NCBI txid 2901164) genome mining, 283 genomes were downloaded from the NCBI GenBank database (accessed June 6th, 2025) (**Table S2**). A preliminary search revealed 1,050 genomes deposited in this database. However, after applying quality criteria (assembly level [>scaffold], exclusion of atypical genomes and those containing 0% completeness or contamination), 283 genomes remained. BLASTp was used for genome mining of potential plastic-degrading enzymes using the following criteria: query coverage > 70% and e-value > 1e-5, according to Carr and colleagues (2022), with modifications.

### *The* Stutzerimonas *pangenome*

The pangenome of 284 *Stutzerimonas* genomes was calculated using Panaroo (v1.5.2) (Tonkin-Hill et al., 2020) using the “--remove-invalid-genes”, “--clean-mode strict”, “--alignment core” and “--core_threshold 0.95” options. The pangenome rarefaction curve was constructed using the “gene_presence_absence” output. Briefly, the output was imported into RStudio (v2024.04.2+764) using the tidyverse library, and a presence/absence gene matrix was generated with the pivot_wider() function. For each sample size (from one to 284 genomes), 20 random replicates of genome subsets were generated. For each subset, the number of core and pangenome genes was calculated, and the results were visualised using the R ggplot2 package (Wickham, 2016). Genes were further classified into cloud genes (0% ≤ strains < 15%), shell genes (15% ≤ strains < 95%), soft core genes (95% ≤ strains < 99%), and core genes (99% ≤ strains ≤ 100%). To understand whether gene content could explain species variation, a principal coordinate analysis (PCoA) was performed using the Panaroo “gene_presence_absence” output file. The gene presence/absence matrix was transposed and binarised to represent gene presence (1) and absence (0). Genome identifiers were standardised to match the metadata downloaded from the NCBI GenBank database. Pairwise Jaccard distances were computed using the vegdist() function (vegan package) and PCoA was performed on the Jaccard distance matrix using the pcoa() function (ape package). The results were again plotted with ggplot2. Homologous sequences to the reference poly(ethylene terephthalate) hydrolase (PETase) protein WP_013984362.1 were identified and redundancy was reduced using CD-HIT with a 100% identity threshold (-c 100) (Limin et al., 2012). The non-redundant homologous protein sequences were aligned using MAFFT (v7) (Katoh et al., 2013) with default parameters. A phylogenetic tree was constructed from the multiple sequence alignment using the Neighbor-Joining method. The tree was visualised and annotated to highlight sequences of interest with iTOL (v6) (Letunic et al., 2024).

## Results

### *Genome of* Stutzerimonas frequens *strain VG-9*

The draft genome sequence of *S. frequens* strain VG-9 was successfully obtained with high quality (**Fig. 1A**). A total of 3,482,256 raw reads were generated, which were processed by removing adapter sequences and trimming the first 15 low-quality bases. Further quality control was performed using a sliding window to discard bases with a quality score below Q20, retaining only sequences with a minimum length of 40 base pairs. The processed reads were assembled into a draft genome spanning 4.68 Mb (accession number: SAMN49487720). The assembly resulted in 60 scaffolds with an N50 of 180 kbp and an average sequencing coverage of 105x (**Table 2**). Annotation with Bakta identified 4,392 protein-coding genes. Quality assessment indicated high completeness, with CheckM estimating 99.03% and BUSCO confirming 99.6%. Furthermore, an ANI value of 96.5% supports the identification of strain VG-9 as *S. frequens* (**Table 2**; **Fig. 1B**).

**Fig. 1.**
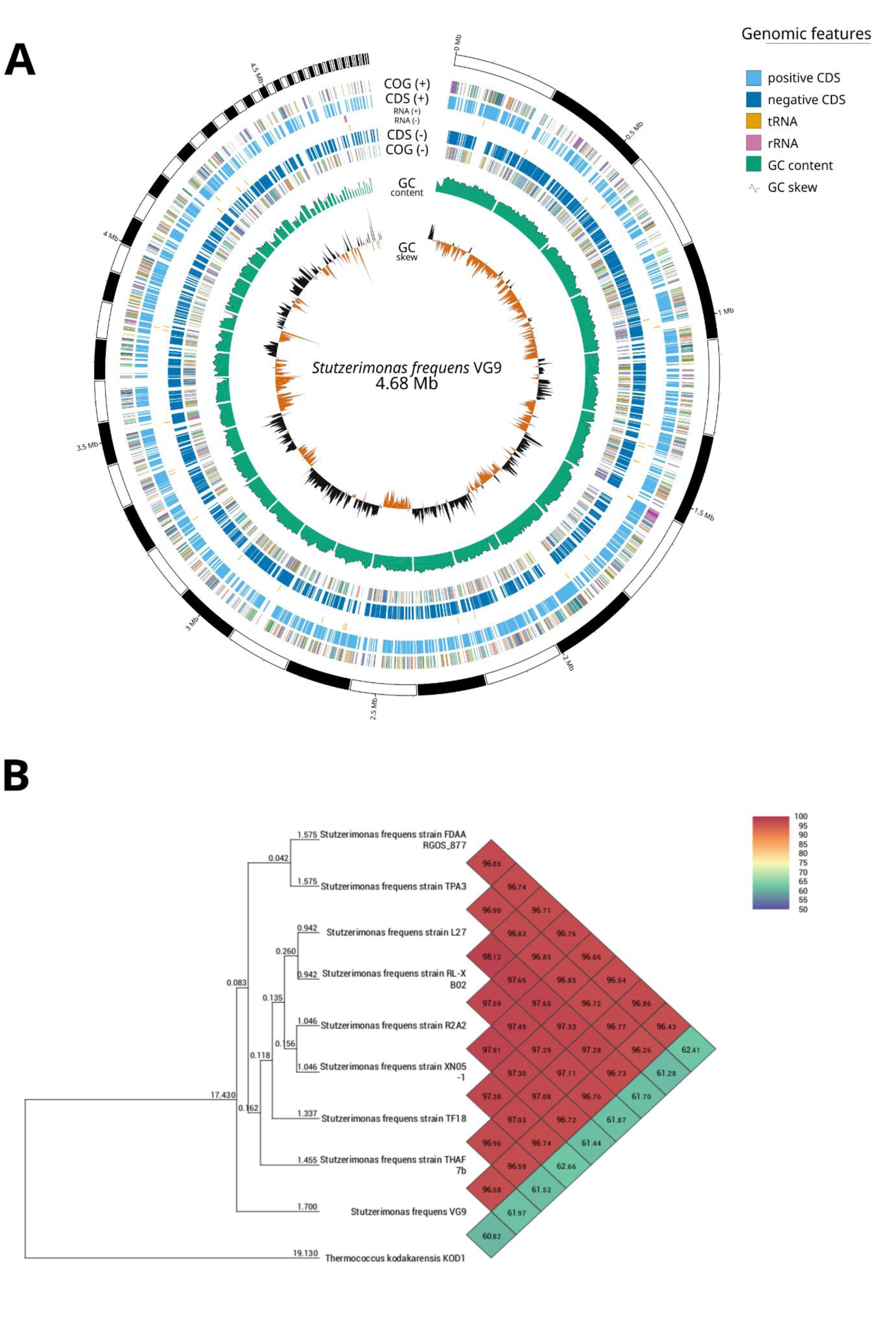
Genomic features and similarity of *Stutzerimonas frequens* VG-9. (**A**) Circular genome map of *S. frequens* VG-9. From the outermost to innermost ring: i) genome size scale; ii) COG functional categories of CDS on the forward strand; iii) coding sequences (CDS) on the forward strand; iv) RNA genes (rRNA, tRNA) on the forward strand; v) RNA genes on the reverse strand; vi) CDS on the reverse strand; vii) COG functional categories of CDS on the reverse strand; viii) GC content; ix) GC skew. (**B**) Average Nucleotide Identity (ANI) heatmap. The matrix shows pairwise ANI values between *S. frequens* VG-9 and eight other complete *S. frequens* genomes, with *Thermococcus kodakarensis* KOD1 included as an outgroup.

The genome was found to harbour three tailed prophage regions, two of which were intact and one incomplete. The first intact phage, PHAGE_Pectob_ZF40_NC_019522(19), is 35.4 kb in length, has a GC content of 62.5%, and encodes 29 proteins, encompassing tail, plate, capsid, terminase, portal, head, and integrase. The second, PHAGE_Pseudo_Dobby_NC_048109(23), spans 46.3 kb, has a GC content of 64.5%, and encodes 48 proteins, including integrase, plate, tail, virion, lysis, head, terminase, and capsid. The incomplete phage, PHAGE_Pseudo_PS_1_NC_029066(7), spans 13.1 kb and has a GC content of 59.5%, harbouring only a tail and an integrase.

The strain also harbours a 27.9 kb plasmid with 99.9% identity and 84% coverage with the p1_PM101005 from *S. stutzeri* strain PM101005 (NZ_CP046903.1). In contrast to the 265 kb complete reference plasmid p1-PM101005, the detected segment most likely represents a partial linear sequence. This may be due to the sequencing method used, which did not enable the yield of a complete circular sequence. Nonetheless, functional annotation of this region revealed the presence of genes associated with plasmid maintenance, including a replication initiation protein (RepA), a partition protein of the ParB family, and an ATPase involved in segregation processes (Soj). These genes are distributed along the contig and are characteristic of extrachromosomal replicons. It also harbours genes associated with mercury resistance, DNA repair (e.g. DNA polymerase, UmuC subunit of DNA polymerase V and UvrD helicase), and several mobile elements, including a family Tn*3* transposase, multiple integrases, and genes encoding phage proteins. Genes corresponding to a toxin–antitoxin system were identified, including a toxic component belonging to the PIN family, as well as associated transcriptional regulators such as XRE and ribbon–helix–helix family proteins (**Fig. S1**).

The genome of *S. frequens* VG-9 harboured genes for several multidrug efflux systems. Four genes encoding components of the membrane multidrug exporter complex MexAB-OprM were identified with high confidence (> 98% coverage and > 80% identity): *mexB* (AAA74437.1), *mexF* (AE004091.2), *mexK* (AE004091.2), and *mexW* (NC_002516.2) (Li et al., 1995). Further, screening with ABRicate revealed seven additional efflux pump genes with lower sequence similarity (coverage and identity < 80%). Three of these were associated with the MexEF-OprN operon and four with the MexJK-OprM operon (Chuanchuen et al., 2002; Fetar et al., 2011). Analysis against the Virulence Factor Database (VFDB) using ABRicate identified 28 virulence-associated sequences. These included 14 genes related to flagellar assembly (*fleN, fleQ, flgG, flgI, flhA, fliA, fliE, fliG, fliI, fliM, fliN, fliP, fliQ, motC*), five involved in twitching motility (*pilG, pilH, pilR, pilT, pilU*), five components of the alginate pathway (*algA, algB, algC, algR, algU*), two components of the Xcp secretion system (*xcpR, xcpT*), and two enzymes (*waaF, waaG*) critical for constructing the lipopolysaccharide (LPS) core region (**Table 1**).

**Table 1.**
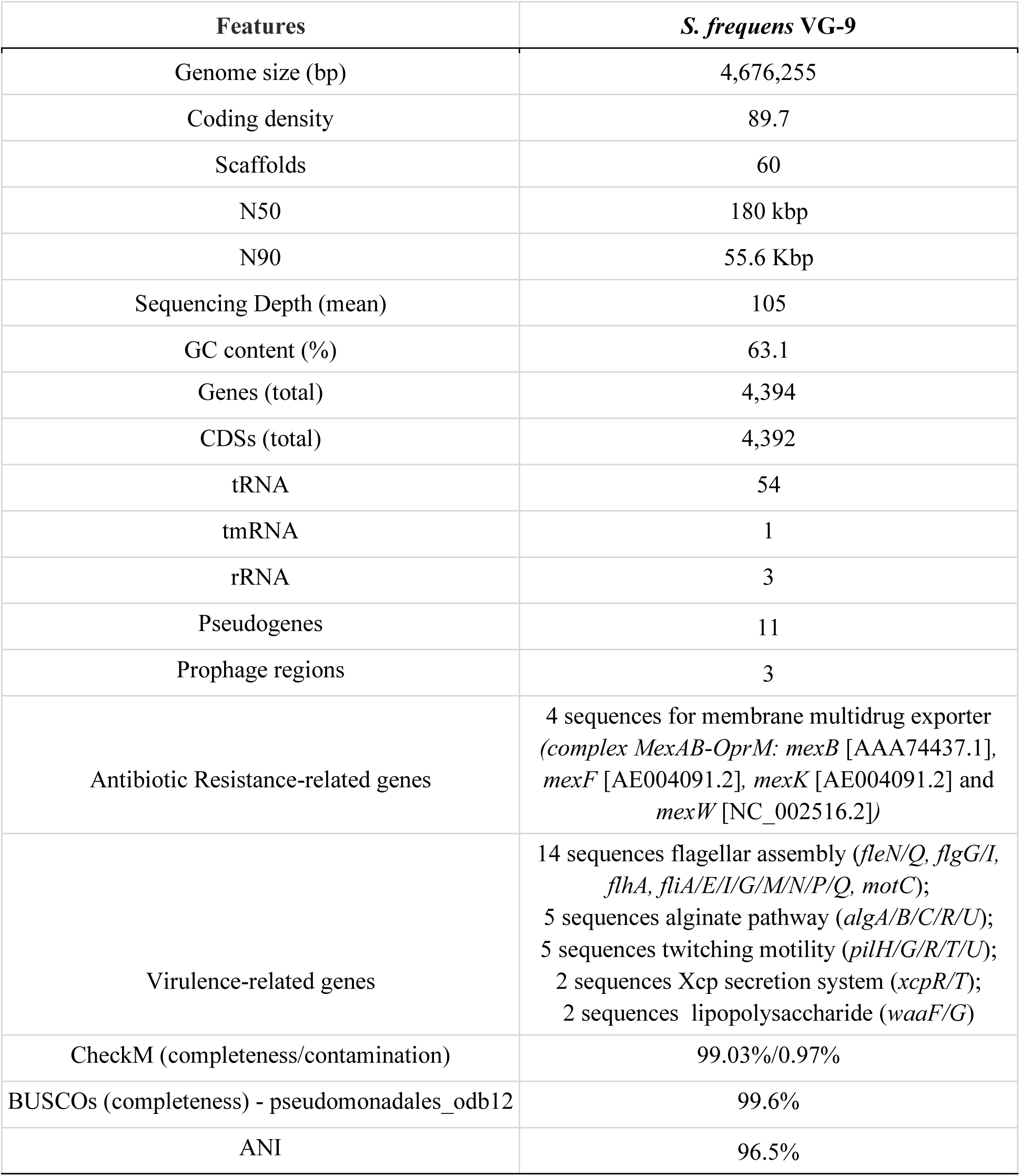
Genomic features of *S. frequens* VG-9.

Genomic annotation via BlastKOALA revealed that *S. frequens* VG-9 encodes a diverse repertoire of secretion systems, dominated by Type VI Secretion System (T6SS) (Coulthurst et al., 2019). This system includes the core structural apparatus (Imp/Vas proteins), the hallmark secreted components Hcp and VgrG, the ATPase ClpV for powering the system, and key regulators like the serine/threonine-protein kinases PpkA and Stk1. Coupled with essential export systems like the Sec and Tat pathways, this secretory capability suggests a high degree of adaptability.

Nine regions encoding biosynthetic gene clusters (BGCs) were detected using antiSMASH, including ribosomally synthesised and post-translationally modified peptides (RiPP)-like (**Fig. 2A**), NI-siderophore-terpene-precursor, redox-cofactor, NAGGN (**Fig. 2B**), terpene, arylpolyene, betalactone, NRPS-like-ectoine. Among the clusters exhibiting high similarity confidence, those for carotenoid and ectoine biosynthesis were detected. In contrast, clusters for the biosynthesis of putrebactin/avaroferrin, aryl polyene (APEVf), and corynecin III/corynecin I/corynecin II exhibited low confidence scores for similarity.

**Fig. 2.**
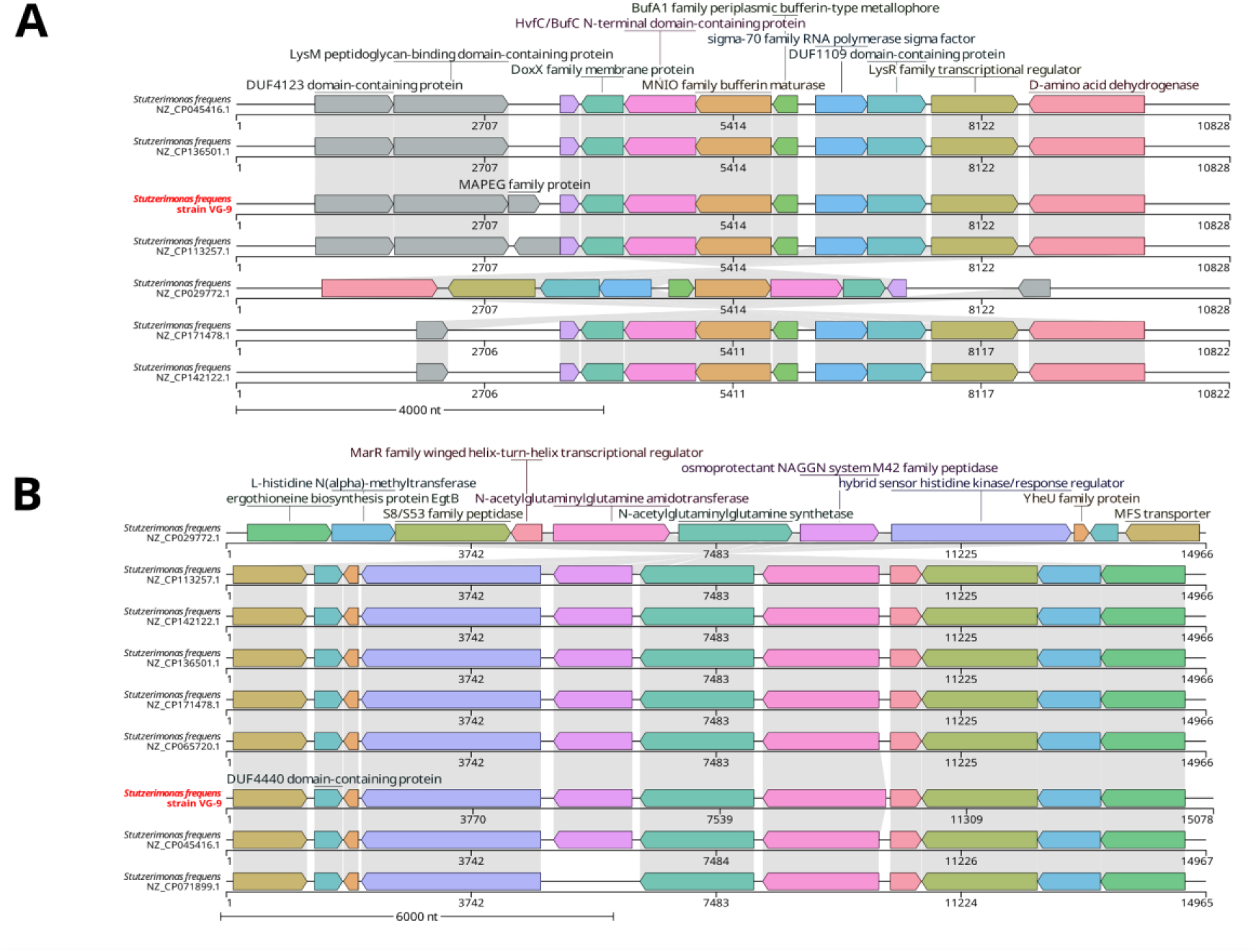
Genomic organization of secondary metabolite biosynthetic gene clusters (BGCs) in *S. frequens* strains. (**A**) RiPP-like and (**B**) NAGGN BGCs are shown. A multiple sequence alignment was performed using *S. frequens* strain FDAARGOS_877 as a reference. The analysis included all *S. frequens* genomes from the Average Nucleotide Identity (ANI) dataset (see Methods). Only strains harbouring the RiPP-like and NAGGN BGCs, as predicted by antiSMASH, are displayed.

### PETase activity in Stutzerimonas: the case of S. frequens VG-9

To assess potential PETase and esterase activity associated with *S. frequens* VG-9, enzymatic activity was evaluated on solid and liquid growth media with several substrates. *S. frequens* VG-9 showed the capacity to hydrolyse Tween 20 and polycaprolactone, while no activity was detected towards Impranil, tributyrin, or BHET in solid media (**Fig. S2**). *In silico* PETase bioprospection revealed that the strain harbours one enzyme with 81,395% identity with PmC with a sequence coverage of 91%. The EC number inferred for the amino acid sequence was EC 3.1.1.101 with high confidence, which refers to the poly(ethylene terephthalate) hydrolases. According to genome annotation, *S. frequens* VG-9 also harbours a MHETase, which was assigned to EC 3.1.1.102 mono(ethylene terephthalate) hydrolases, with low confidence.

To further identify the potential degradation of PET dimers by strain VG-9, cultures with BHET and MHET as carbon sources were carried out (**Fig. 3A**). Complete consumption of BHET occurred in the first 48 hours of cultivation in LB medium in the presence of this dimer (**Fig. 3B**). Putative MHET formation was observed, although peak retention times were slightly different when compared to the reference curves (**Fig. S3**). This may be due to the formation of BHET oligomers or an intermediate compound between BHET and MHET, from which TPA was subsequently released over the course of cultivation. Different concentrations of salts and other biomolecules between the samples and the standards may also exert an influence and further characterisation is needed to pinpoint what exactly was generated. However, in the context of the analysis, MHET remains the most plausible candidate. On the other hand, the highest bioconversion efficiency was obtained using MHET as a carbon source, with 66.3% conversion and productivity approximately 2.8 times higher than in the culture with BHET after 72 h of cultivation. Thus, it was necessary to extend the cultivation time supplemented with BHET to increase TPA formation. Then, after 20 days of cultivation with BHET, it was possible to obtain 342.9 mg.L^-1^ of TPA and a conversion of 70.6%. These results are consistent with the genomic identification of both PETase and MHETase, indicating that BHET hydrolysis by the wild-type strain may be limited under the tested conditions and that MHETase is likely more active, enabling conversion of MHET to TPA.

**Fig. 3.**
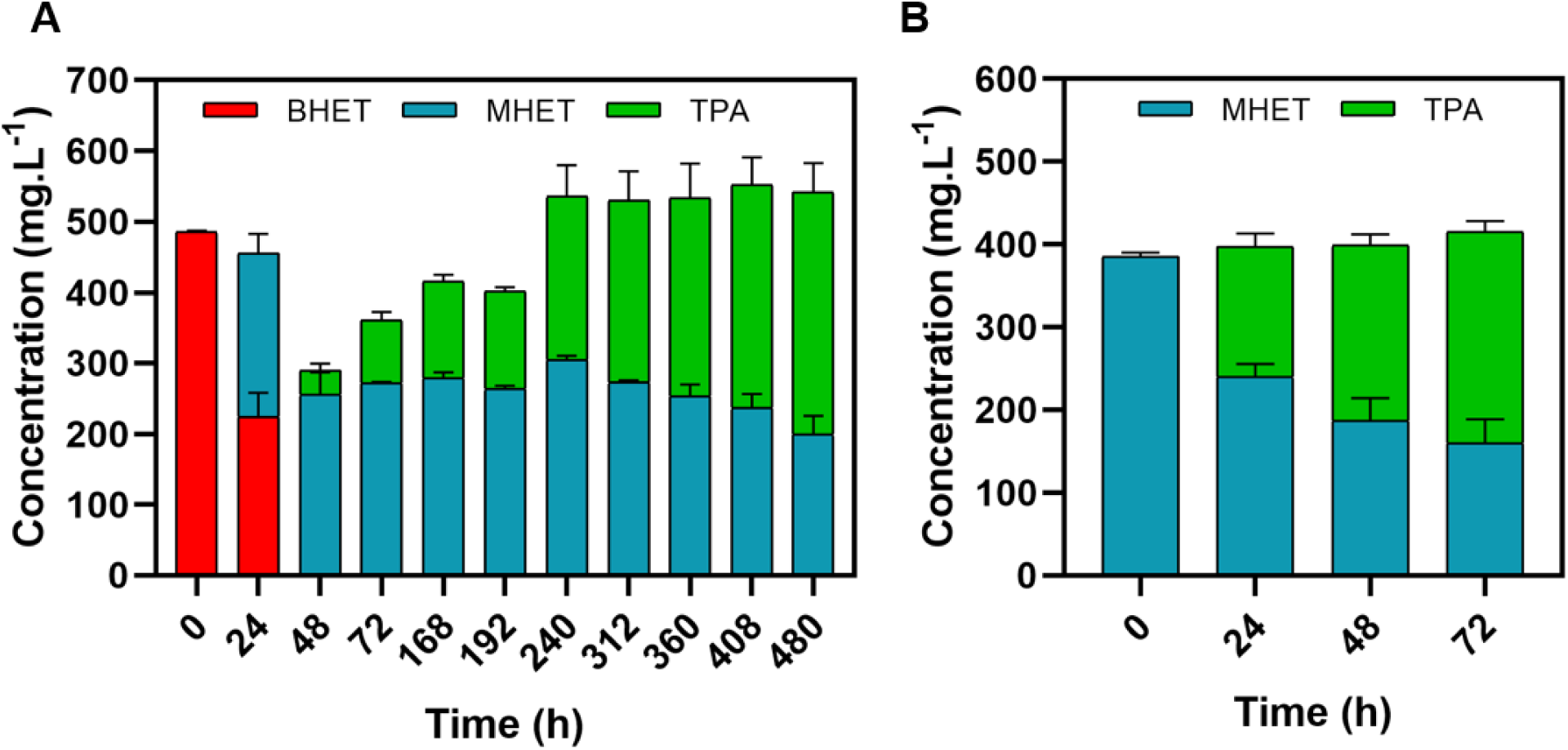
Degradation profile of PET dimers by *S. frequens* VG-9 cultivated in LB medium supplemented with BHET, shown in red, (A) and MHET, shown in dark green (B). Panel A shows the progressive conversion of BHET into MHET followed by the accumulation of terephthalic acid (TPA, shown in light green) over time. Panel B illustrates MHET degradation with a concomitant increase in TPA concentration. Data represent mean ± standard deviation from duplicate experiments.

### PETase activity in Stuzterimonas: genomic insights

A total of 284 *Stutzerimonas* genomes were annotated and investigated for the presence of PETase homologs. From these, 86 (30.3%) were identified as *Stutzerimonas stutzeri*, 81 (28.5%) as *Stutzerimonas nitrititolerans*, 46 (16.2%) as *Stutzerimonas kunmingensis*, 22 (7.7%) as *Stutzerimonas balearica*, 16 (5.6%) as *Stutzerimonas frequens*, eight (2.8%) as *Stutzerimonas xanthomarina*, seven (2.5%) as *Stutzerimonas chloritidismutans*, five (1.7%) as *Stutzerimonas* sp., four (1.4%) as *Stutzerimonas zhaodongensis*, three (1.1%) as *Stutzerimonas degradans*, and two as *Stutzerimonas marianensis* (0.7%). The species *Stutzerimonas decontaminans*, *Stutzerimonas azotifigens*, *Stutzerimonas tarimensis*, and *Stutzerimonas urumqiensis* each were represented by one genome, totalling 1.4% of the total. The genome of a *S. frequens* strain isolated by our research group was included in the analysis (accession number: SAMN49487720).

From the downloaded genomes, most were deposited in the form of scaffolds (*n* = 220, 78%), followed by complete genomes (*n* = 60, 21%) and chromosomes (*n* = 3, 1%). Genome sizes ranged from 3,589,459 to 5,821,725 bp (average = 4,471,810 ± 294,787 bp), while GC content ranged from 59.5% to 67% (**Fig. S4**). From the available metadata, the isolation sources were varied, with a significant proportion (*n* = 114, 40.1%) associated with “urban” environments, 17.60% (*n* = 50) being classified as “unknown”, 9.5% (*n* = 27) with marine environments, and 9.2% (*n* = 26) with hospital settings, among others (**Table S1**).

A total of 743 PETase homology hits (identity ≥ 65%; sequence coverage > 70%) were detected across the *Stutzerimonas* genomes, using a custom PAZY-derived database of 75 PETase homologues (**Table S1**) against 284 *Stutzerimonas* genomes (**Table S2**). Of these, the most detected enzyme was PpEst from *Pseudomonas pseudoalcaligenes* (285 hits, 38.3%), followed by EstB from *Pseudomonas* sp. (283 hits, 38.1%), and PmC from *Pseudomonas mendocina* ATCC 53552 (146 hits, 19.6%) (**Fig. 4A**). *S. zhaodongensis* (GCF 047758595.1) was the genome with the highest number of hits, with four potential genes encoding PET-active enzymes (WP_122164697.1, WP_255830209.1, WP_415845153.1, and WP_415846081.1). *S. stutzeri*, *S. nitrititolerans*, and *S. kunmingensis* stood out for the highest number of hits, while *S. zhaodongensis* displayed the highest diversity of hits. The EstB (WP_085690612.1) enzyme was detected across all *Stutzerimonas* species, while the PpEst enzyme was detected in all species except for *S. tarimensis* and *S. urumqiensis* (**Fig. 4B**).

**Fig. 4.**
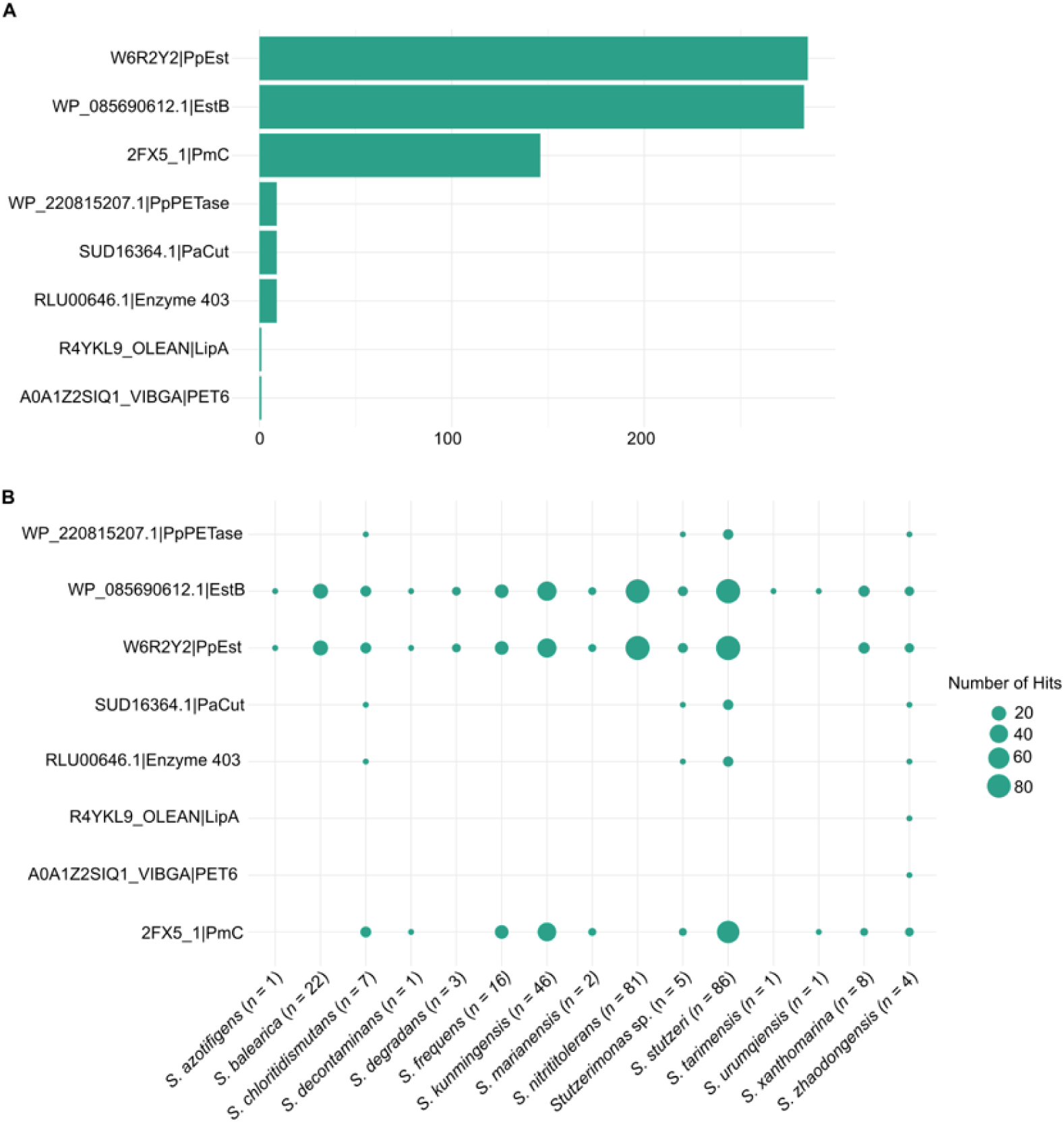
Overview of mining PETase homologs in *Stutzerimonas* genomes. (A) Number of times each enzyme has been detected in the genomes (see **Table S1** for accession number/enzyme name). (B) Distribution of PETase homologs identified in 284 *Stutzerimonas* genomes. Bubbles represent the hits (accession number|enzyme) per species, with sizes proportional to the number of hits detected.

### Stutzerimonas *pangenome*

Pangenome analysis of 284 *Stutzerimonas* genomes (283 public and one newly sequenced *S. frequens* VG-9) reconstructed a total gene repertoire of 35,045 genes. These genes were categorized based on their frequency across strains into 86 core genes (99% ≤ strains ≤ 100%), 1,163 soft core genes (95% ≤ strains < 99%), 3,994 shell genes (15% ≤ strains < 95%), and 29,082 cloud genes (0% ≤ strains < 15%) (**Fig. 5A**). The pangenome rarefaction curve demonstrated a persistently open profile, indicating that the total gene repertoire continues to expand with the inclusion of additional genomes without reaching saturation (**Fig. 5B**).

**Fig. 5.**
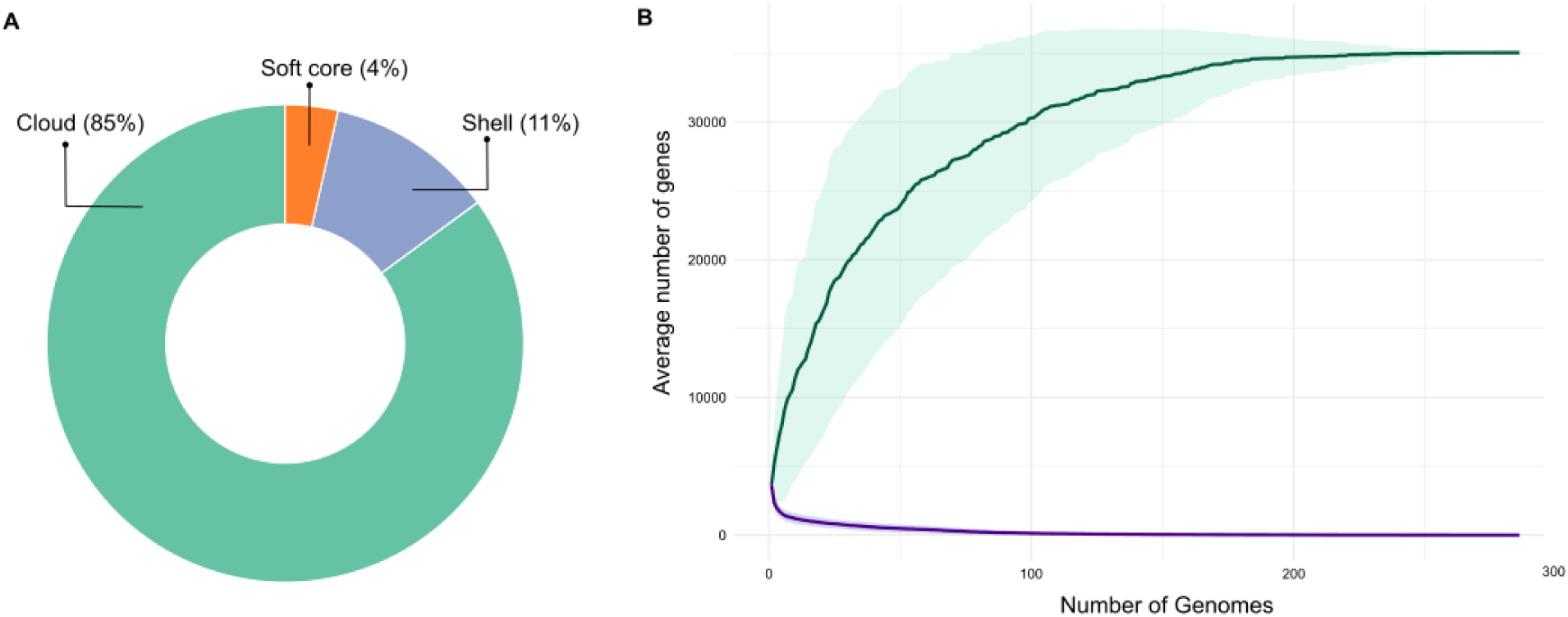
Overview of the *Stutzerimonas* pangenome. (A) Distribution of core, soft core, shell and cloud genes across 284 *Stutzerimonas* genomes. (B) Rarefaction curves of the pangenome (green) and core genome (purple). The curves represent the mean values from 20 random replicates, with shaded areas indicating the standard deviation.

Analysis of the pangenome distribution of PETase homologs showed that key candidates reside in the accessory genome. The ‘poly(ethylene terephthalate) hydrolase family protein’ gene (WP_013984362.1), for instance, was present in 146 genomes (∼51.4%), primarily in species such as *S. stutzeri* and *S. frequens*. Homologous sequences to the reference poly(ethylene terephthalate) hydrolase (PETase) protein WP_013984362.1 were identified and redundancy was reduced using CD-HIT with a 100% identity threshold (-c 100), resulting in a non-redundant set of 56 protein sequences. This prevalence level places it within the shell gene category, highlighting its role as a flexible genomic trait shared by a significant subset of the genus (**Figs. 6** and **7**).

**Fig. 6.**
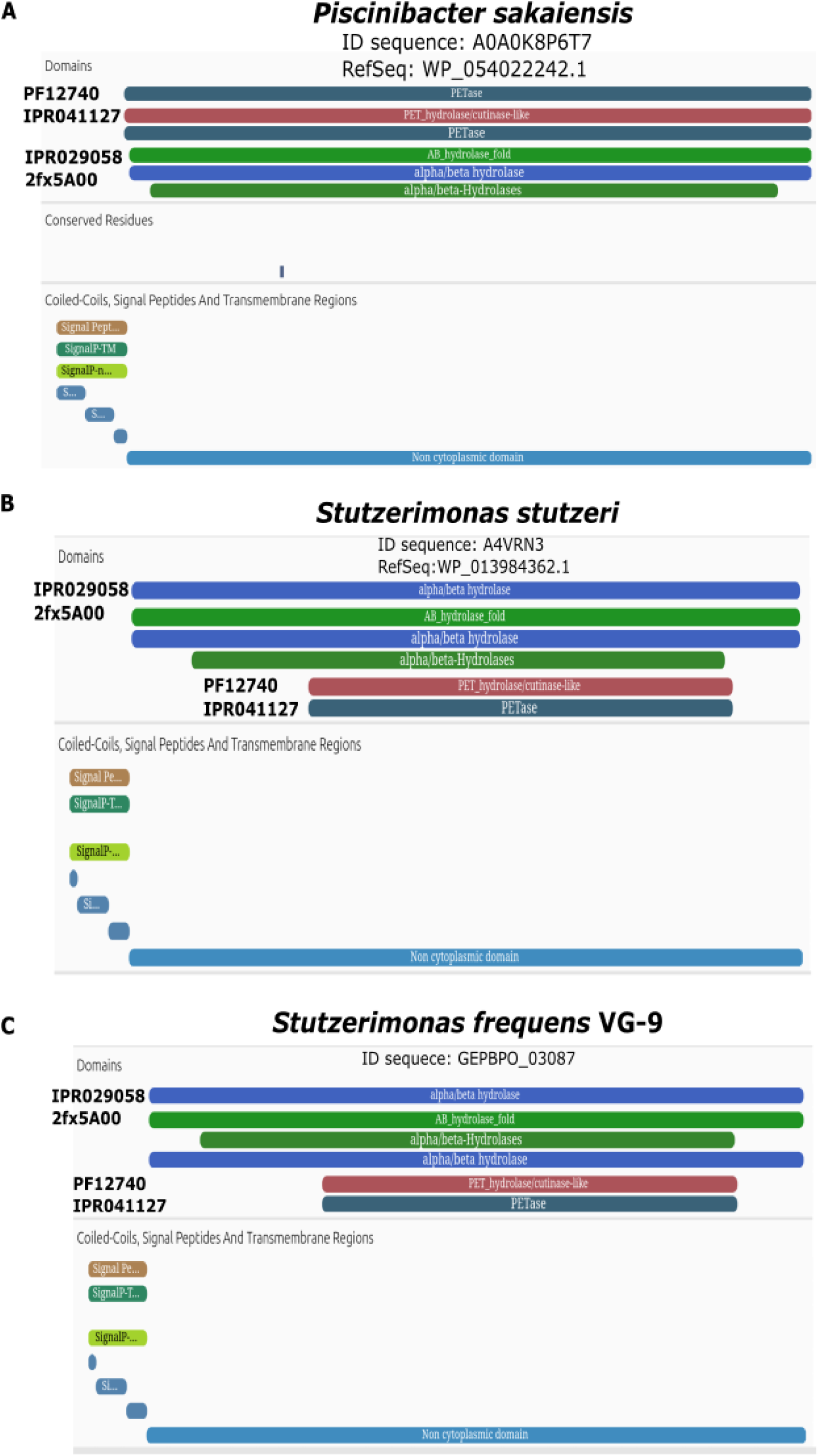
Structural analysis of the poly(ethylene terephthalate) hydrolase family protein across *Stutzerimonas* genomes. (**A-C**) Domain architecture. Representative sequences from *Piscinibacter sakaiensis* (WP_054022242.1), *S. stutzeri* (WP_013984362.1), and *S. frequens* VG-9 (GEPBPO_03087) all contain the conserved domains IPR029058 (model: 2fx5A00), Pfam PF12740, and IPR041127. A signal peptide is predicted in the N-terminal region (amino acids 23-27) of each sequence.

**Fig. 7.**
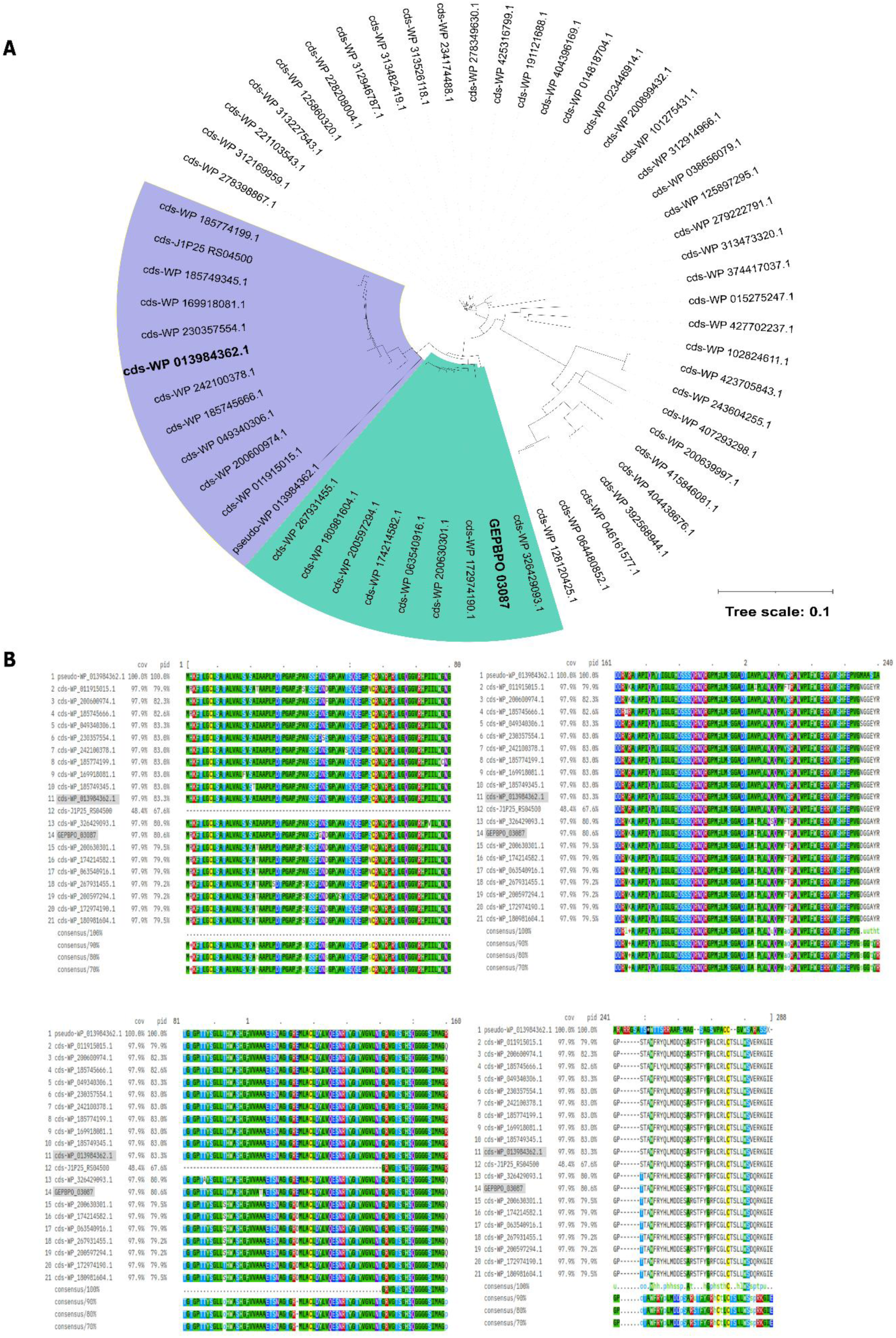
Evolutionary analysis of the poly(ethylene terephthalate) hydrolase family protein across *Stutzerimonas* genomes. (**A**) Neighbor-Joining phylogenetic tree based on protein sequence alignment. The tree was constructed from an alignment of 58 non-redundant homologous protein sequences (289 aa each) using MAFFT (v7). The reference sequence from *S. stutzeri* (WP_013984362.1) is highlighted in purple, and the sequence from *S. frequens* VG-9 (GEPBPO_03087) is highlighted in green. (**B**) Multiple Sequence Alignment (MSA). A segment of the alignment is shown, illustrating the conserved regions and sequence variation among the homologous protein sequences from the analysed *Stutzerimonas* genomes. Scale bar represents the number of amino acid substitutions per site.

## Discussion

The growing problem of plastic pollution has intensified the search for new biodegradation strategies, with microbial enzymes emerging as promising candidates. Genome mining approaches have emerged as powerful tools for identifying candidate biocatalysts within publicly available genomic data. However, the functional relevance of these candidates often remains to be experimentally demonstrated (Ali et al., 2021; Zhu et al., 2022; Dhali et al., 2024). Members of the Pseudomonadaceae family, in particular, have shown a remarkable propensity for plastic degradation, consistent with the presence of diverse hydrolytic enzymes that may, under specific conditions, act on synthetic polymers (Bollinger et al., 2020; Sagong et al., 2021; Han et al., 2024). This potential motivated the investigation into the recently established genus *Stutzerimonas*, a taxon split from *Pseudomonas* in 2022 based on distinct phylogenetic markers (Lalucat et al., 2022). While the plastic-degrading ability of the reclassified *Stutzerimonas stutzeri* had been previously noted (Howard et al., 2022; Li et al., 2025), the potential of the broader genus remained largely unexplored.

Our investigation into plastic-degrading potential across Pseudomonadaceae began with a broad genomic screen of 100 publicly available genomes (**Appendix A**). This preliminary analysis revealed a striking concentration of PETase homologs within the *Halopseudomonas* genus, which accounted for all 18 identified hits. This finding corroborates growing evidence of *Halopseudomonas* proficiency in degrading various polyesters, underscoring their genetic versatility despite possessing compact genomes (∼ 4 Mbp). Notably, this screen also highlighted a methodological consideration: the conserved α/β-hydrolase fold architecture of many PET-degrading enzymes can produce multiple database matches for individual genes, emphasising the need for rigorous sequence curation to ensure accurate annotation.

A more directed approach was carried out with the functional characterisation of a newly isolated strain, *S. frequens* VG-9, which demonstrated the ability to hydrolyse the model polyester PCL and Tween 20 (Almeida et al., 2019). Even though no activity towards BHET was observed on solid media, liquid cultures supplemented with BHET and MHET indicated the degradation of these substrates and the formation of TPA over time. While these observations indicate hydrolytic activity toward soluble PET-derived intermediates, they do not necessarily demonstrate efficient PET depolymerization, as similar profiles may arise from nonspecific esterases, co-metabolic processes, or intracellular hydrolysis following substrate uptake. Nonetheless, this phenotypic activity motivated whole-genome sequencing to uncover potential genetic determinants underlying this behaviour. The genomic characterisation of VG-9 revealed a repertoire of features that underscore its adaptability and potential ecological fitness in complex microbial environments. Notably, a Type VI Secretion System (T6SS) was identified, complete with structural components and regulators such as PpkA and Stk1 (Silverman et al., 2012; Flaugnatti et al., 2025; Pei et al. 2025). This system suggests the capacity for interbacterial competition, an advantageous trait in the complex microbial consortia found in plastic-polluted environments. Furthermore, the genome harbours multiple biosynthetic gene clusters (BGCs) for bioactive metabolites, including those for carotenoids and ectoine, which likely confer stress tolerance under the oligotrophic and potentially toxic conditions of plastic degradation niches (Tong et al., 2024). The genome also features BGCs for antimicrobial peptides, including non-ribosomal peptide synthetases (NRPS) and RiPPs, suggesting a capacity for interbacterial competition (Baltz et al., 2021).

The genomic resilience of *S. frequens* VG-9 is further highlighted by the presence of intrinsic antibiotic resistance genes (e.g., *mexB*, *mexF*, *rsmA*, *adeF*), consistent with reports for other environmental *Stutzerimonas* strains (Hussain et al., 2023; Devkar et al., 2025). These genes are likely part of core regulatory mechanisms rather than acquired resistance, but their presence reinforces the strain’s hardy genetic background. Crucially, the combination of robust secretory capabilities (T6SS, Sec, Tat pathways) and stress adaptation mechanisms highlights genomic features that may support the survival and activity of hydrolytic enzymes under environmentally challenging conditions.

Building on the insights from *S. frequens* VG-9, we expanded our scope to conduct a comprehensive *in silico* mining of PETase homologs across the entire genus. Two hundred and eighty-three high-quality genomes were retrieved from NCBI (accessed June 2025), supplementing them with the newly sequenced genome of the PCL-degrading *S. frequens* VG-9. This strategy allowed us to move from a single-strain analysis to a genus-wide assessment, systematically evaluating the distribution and diversity of potential plastic-degrading enzymes in this underexplored bacterial group. Our comprehensive genome mining across 284 *Stutzerimonas* genomes revealed a significant reservoir of candidate hydrolases related to known PET-active enzymes, identifying 743 high-confidence homologs distributed across the genus. The enzymes PpEst (38.3%) and EstB (38.1%) were the most prevalent, followed by PmC (19.6%), indicating a conserved hydrolytic machinery within this taxonomic group. The near-ubiquitous distribution of EstB across all examined species suggests it may represent an ancestral hydrolase, maintained due to its broad substrate specificity for natural polyesters, which fortuitously pre-adapts it to degrade synthetic counterparts (Wagner et al., 2009).

The distribution of these homologs was not uniform, revealing distinct ecological and genomic patterns. Notably, *S. stutzeri*, *S. nitrititolerans*, and *S. kunmingensis* accounted for the highest number of hits, whereas *S. zhaodongensis* (GCF_047758595.1) exhibited the greatest genetic diversity, harbouring four distinct PETase-like genes (WP_122164697.1, WP_255830209.1, WP_415845153.1, WP_415846081.1). This genomic redundancy suggests potential for functional specialisation or synergistic interactions among different hydrolases, a strategic feature that could be harnessed in engineered microbial consortia for enhanced plastic degradation. Ecological metadata further revealed that strains isolated from urban (40.1%) and marine (9.5%) environments possessed the highest diversity of PETase homologs. This distribution strongly implies an environmental selection for plastic-degrading capabilities in these anthropogenically impacted niches, where microbial communities are consistently exposed to plastic waste. However, alternative explanations, such as selection for general stress tolerance or xenobiotic metabolism, cannot be excluded. Critically, these *in silico* predictions were supported by preliminary functional screening, demonstrating the capacity of *S. frequens* VG-9 to hydrolyse polyesters in solid (PCL and Tween 20) and liquid (BHET and MHET) cultures. This phenotypic activity bridges genomic prediction with tangible catabolic activity, suggesting the role of *Stutzerimonas* as a source of plastic-active enzymes and prioritising these candidates for further biochemical characterisation.

Building on these family-wide observations, we conducted the first comprehensive pangenome analysis of the recently reclassified *Stutzerimonas* genus, incorporating 283 public genomes alongside our PCL-degrading isolate *S. frequens* VG-9. The analysis revealed an open pangenome architecture characterised by a small core genome and extensive accessory repertoire, a genomic structure indicative of high adaptability to diverse ecological niches. Crucially, we found that key PETase homologs, including the poly(ethylene terephthalate) hydrolase family protein, reside predominantly in the shell genome (present in ∼51% of strains). This distribution pattern is consistent with the hypothesis that polyester degradation represents a facultative trait, likely disseminated through horizontal gene transfer and maintained by environmental selection.

The limited core genome of *Stutzerimonas* aligns with previous pangenome studies of Pseudomonadaceae, such as *Pseudomonas aeruginosa* (Freschi et al., 2018). Although pangenome analyses within this family have historically focused on pathogenic *Pseudomonas* species, our work demonstrates the importance of extending comparative genomics to understudied relatives. The concentration of plastic degradation potential in specific genera, with *Halopseudomonas* representing a specialised source of abundant homologs and *Stutzerimonas* exhibiting a more sporadic accessory distribution, reveals distinct evolutionary strategies for polymer metabolism within the family.

All in all, this study identifies *Stutzerimonas* as a valuable reservoir of candidate hydrolases for novel enzyme discovery within an open, adaptable pangenome. The phylogenetic framework provided by our analysis enables targeted bioprospecting, while the comparative context with *Halopseudomonas* provides broader insights into the distribution of plastic-degrading potential across Pseudomonadaceae. Our integrated approach, combining *in silico* mining, functional screening, and pangenome analysis, has pinpointed priority candidates, such as those in *S. frequens* VG-9, for subsequent characterization thereby paving the way for their further exploration in the context of biotechnological applications.

## Author contributions

ALBC: Conceptualisation, Data curation, Visualisation, Investigation, Methodology, Formal analysis, Writing – original draft, review & editing. MMO: Data curation, Investigation, Methodology, Formal analysis, Writing – review & editing. YIIR: Data curation, Investigation, Methodology, Formal analysis. BFRO: Conceptualisation, Validation, Writing – review & editing. NFS: Methodology, Formal analysis. MAZC: Methodology, Writing – review & editing. JHW: Funding acquisition, Conceptualisation, Validation, Supervision, Writing – review & editing. MSL: Funding acquisition, Project administration, Resources, Supervision, Writing – review & editing.

## Abbreviations

PET: polyethylene terephthalate
PETase: PET-active enzyme
NCBI: National Center for Biotechnology Information
PCL: polycaprolactone
PAZy: Plastics-Active Enzymes Database

