## Supplementary figures for "Genome Mining and Pangenome Analysis of the *Stutzerimonas* Genus: a Novel Source of Plastic-Degrading Enzymes"

### Supplementary Material

Index Page

**Fig. S1**.......................................................................................................... 2

**Fig. S2**.......................................................................................................... 3

**Fig. S3**.......................................................................................................... 4

**Fig. S4**.......................................................................................................... 5

**Appendix A**................................................................................................... 6

**Fig. S5**.......................................................................................................... 7

**Fig. S6**.......................................................................................................... 8

**References**....................................................................................................9

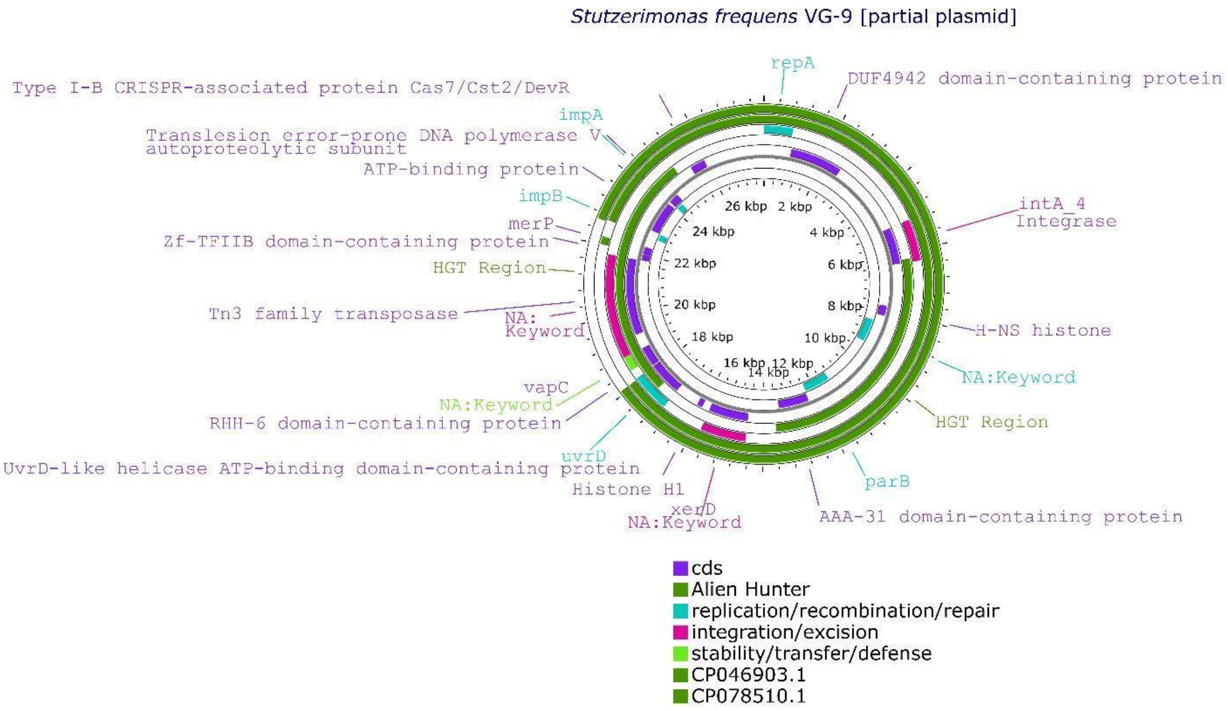


**Fig. S1.** Plasmid map showing the genetic organisation of contig 32, where a partial plasmid was detected in the *Stutzerimonas frequens* VG-9 genome. The distinct features are colored according to the legend.

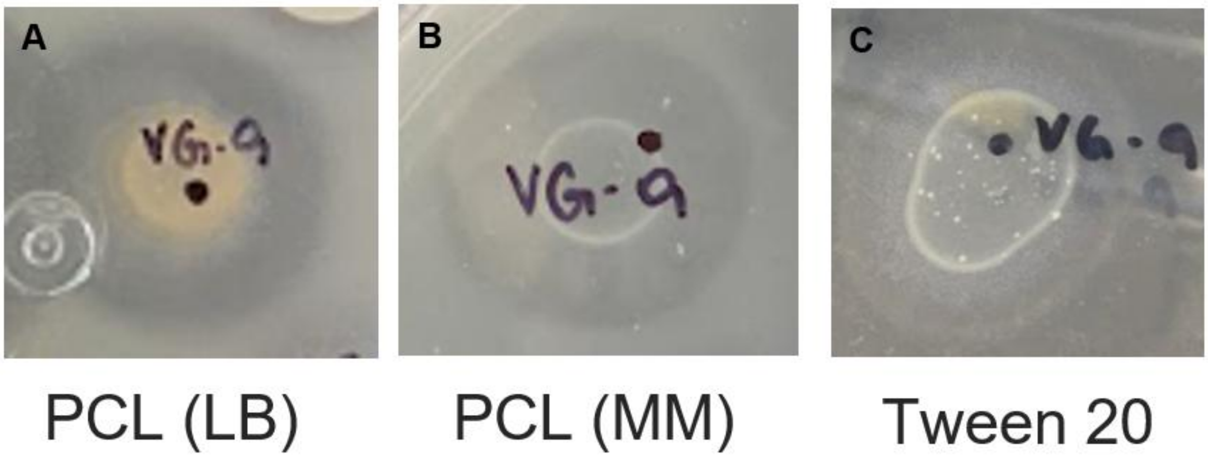


**Fig. S2**. Enzymatic activity of *S. frequens* VG-9 on Lysogenic Broth (LB; A) and minimum medium supplemented with 0.1% PCL (B), as well as minimum medium supplemented with 1% Tween 20 (C).

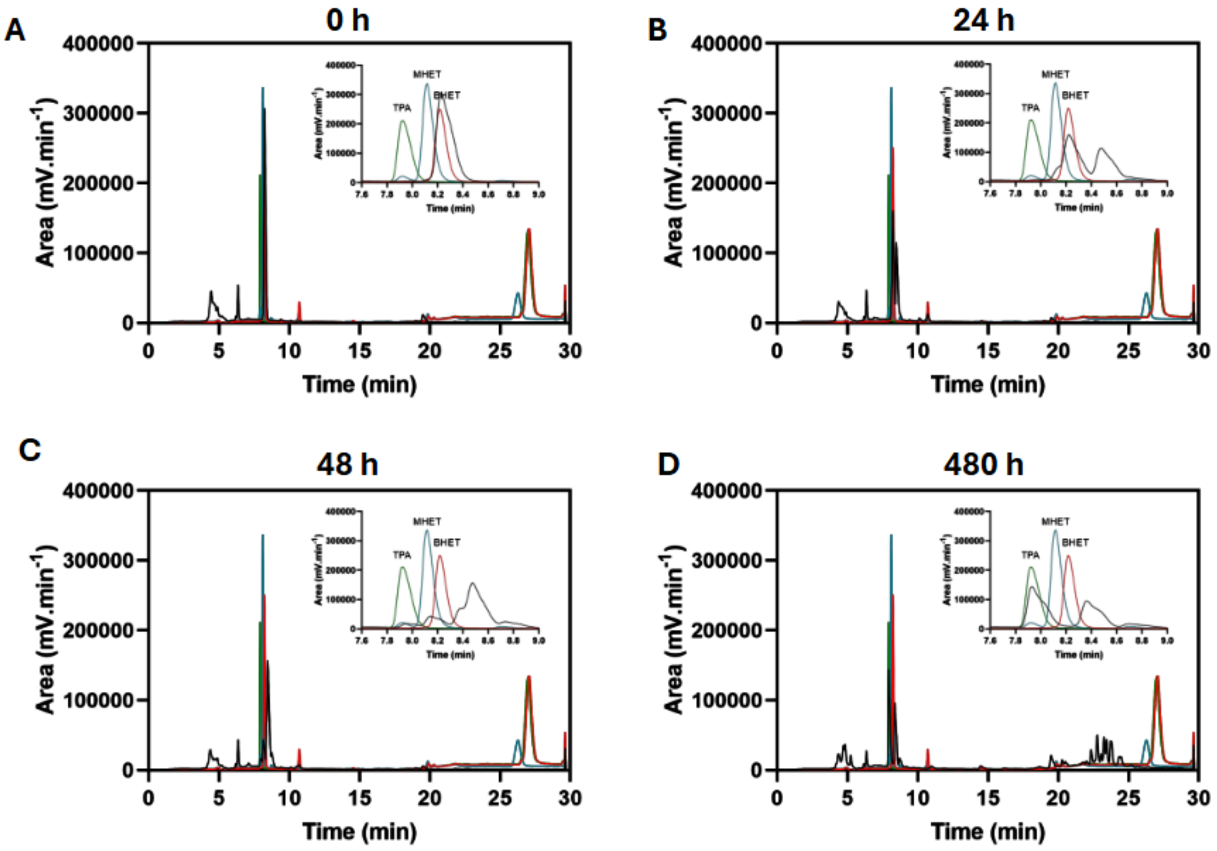


**Fig. S3.** HPLC chromatograms obtained from cultures of *S. frequens* VG-9 (shown in black) in the presence of BHET after 0 (A), 24 (B), 48 (C) and 480 hours (D). In each figure, the retention times between 7 and 9 minutes were zoomed in for better visualisation. Reference compounds TPA, BHET, and MHET are shown green, red, and blue, respectively.

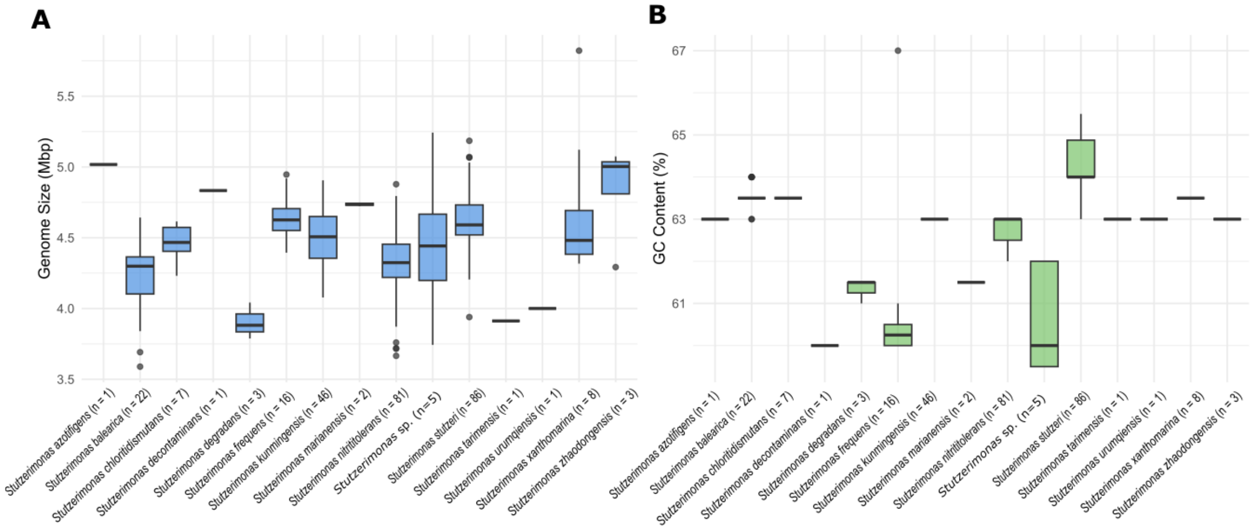


**Fig. S4**. Genome size and GC content of all 284 *Stutzerimonas* genomes, distributed across 14 species and 1 undefined species (sp.). Genome sizes (A) are given in megabase pairs and represented by the blue boxplots. GC content (B) is expressed as a percentage (%) and represented by the green boxplots.

**Appendix A:**

**Genome selection, mining and annotation**

For genome mining of *Pseudomonadaceae*, relevant genomes were obtained from the NCBI Datasets' genome browser (https://www.ncbi.nlm.nih.gov/datasets/genome/, accessed June 6$^{th}$, 2025). A total of 100 publicly available and high-quality genomes from this family were selected randomly and downloaded from the NCBI database in June 2025 (**Table S3**). This random selection aimed to include isolates from diverse environmental sources, ensuring variability and avoiding selection bias. From these, 50 were identified as *Pseudomonas*, 21 as *Halopseudomonas*, 12 as *Stutzerimonas*, ten as *Phytopseudomonas*, and seven as *Azotobacter*. Genome annotation was carried out in a standard approach using Prokka (v. 1.14.6) with default parameters (Seeman, 2014).

Overall, 100 genomes from the Pseudomonadaceae family were downloaded from the NCBI GenBank database (June 6th, 2025), annotated and investigated for the presence of PETase homologs using our customized PETase database. From these, most were deposited as complete genomes ($n$ = 67, 67%), followed by scaffolds ($n$ = 13, 13%), contigs ($n$ = 20, 20%), and chromosomes ($n$ = 2, 2%). Genome sizes ranged from 3,211,846 to 7,352,680 bp (average = 5,400,959 ± 1,011,058 bp), while GC content ranged from 57.5% to 66% (**Fig. S5**).

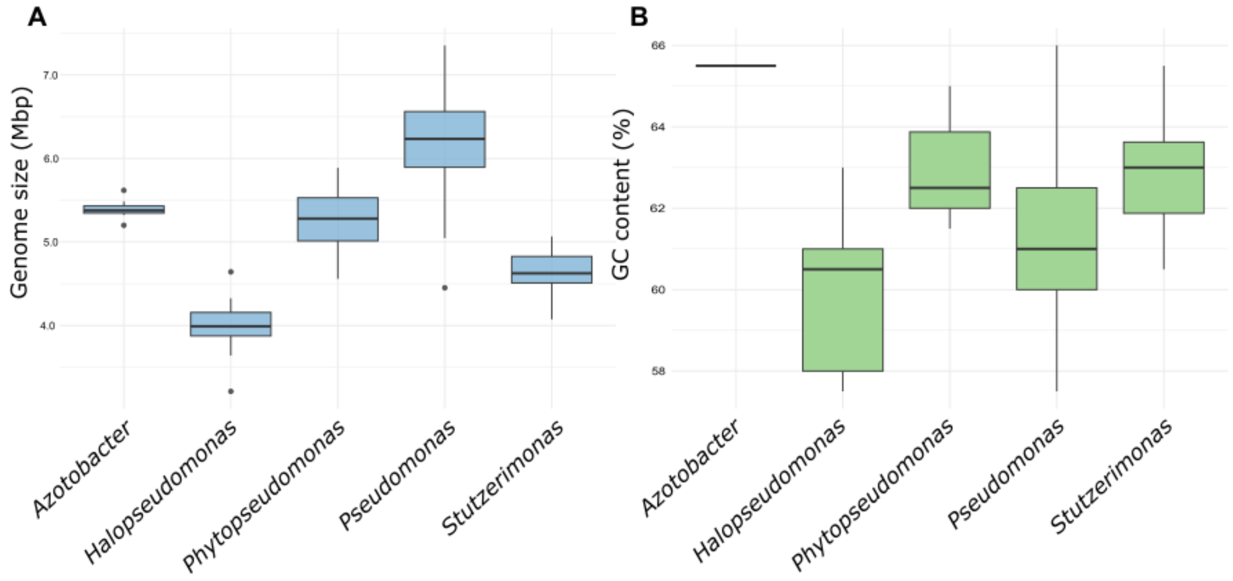


**Fig. S5**. Genome size and GC content of 110 Pseudomonadaceae genomes. Genome sizes (A) are given in megabase pairs and represented by the blue boxplots. GC content (B) is expressed as a percentage and shown in the green boxplots.

A genome mining strategy was employed to investigate the presence of potential PETase homologues in 100 Pseudomonadaceae genomes, rendering 18 hits across 14 genomes, all belonging to the *Halopseudomonas* genus. From these, the Hfor_PE-H enzyme from *Halopseudomonas formosensis* was the most frequently detected (6 hits, 33.3%), followed by PbauzCut from *Halopseudomonas bauzanensis* (4 hits, 22.2%), and PE-H (3 hits, 16.6%) from *Halopseudomonas aestusnigri* VGXO14. The source of isolation of these strains varied, mainly from bottom sediments (**Fig. S6**).

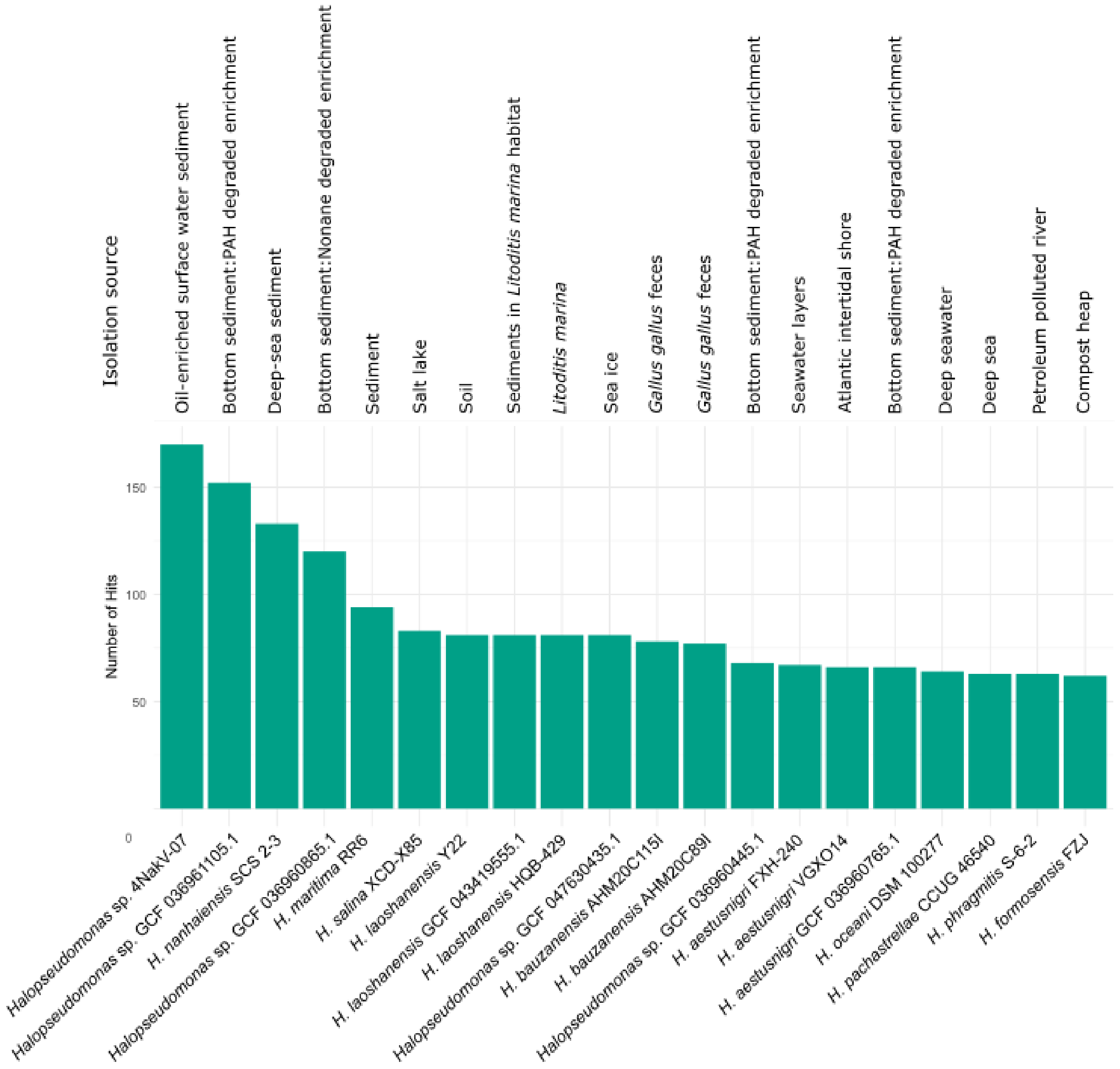


**Fig. S6**. Top 20 Pseudomonadaceae genomes with the highest number of hits with PET-active enzymes and their respective isolation sources.

Our analyses revealed the *Halopseudomonas* genus as a prolific source of novel potential PETases. This is in agreement with previous studies that have delved into the *in vitro* characterization of *Halopseudomonas* isolates capable of degrading polybutylene succinate-co-adipate (PBSA), polycaprolactone (PCL), and polybutylene adipate terephthalate (PBAT), as well as poly(ester-urethane) coatings (de Witt et al., 2023; Soulenthone et al., 2025). Despite its smaller genome size (~ 4 Mb) compared with other members in this family, these bacteria appear to be endowed with the ability to metabolize a range of pollutants and to encode industrially relevant biocatalysts, suggesting high genetic versatility (Kruse et al., 2023). Indeed, in this study, different *Halopseudomonas* species from distinct ecosystems demonstrated the potential to encode multiple putative PETases. The highest number of hits in this genus may be explained by the fact that a single gene often matched multiple enzymes in our database. Many PETases are α/β fold hydrolases that share similar domains and, therefore, may lead to multiple hits. To the best of our knowledge, there is no current bitscore threshold that can be applied to assign a given sequence to a specific family using BLASTp (Punta et al., 2011). Therefore, it is strongly recommended that each hit should be carefully curated in terms of sequence coverage and identity to minimise the risk of false positives.
